# GRAM: A generalized model to predict the molecular effect of a non-coding variant in a cell-type specific manner

**DOI:** 10.1101/482992

**Authors:** Shaoke Lou, Kellie A. Cotter, Tianxiao Li, Jin Liang, Hussein Mohsen, Jason Liu, Jing Zhang, Sandra Cohen, Jinrui Xu, Haiyuan Yu, Mark Rubin, Mark Gerstein

## Abstract

There has been much effort to prioritize genomic variants with respect to their impact on “function”. However, function is often not precisely defined: Sometimes, it is the disease association of a variant; other times, it reflects a molecular effect on transcription or epigenetics. Here we coupled multiple genomic predictors to build GRAM, a generalized model, to predict a well-defined experimental target: the expression-modulating effect of a non-coding variant in a cell-specific manner. As a first step, we performed feature engineering: using a LASSO regularized linear model, we found transcription factor (TF) binding most predictive, especially for TFs that are hubs in the regulatory network; in contrast, evolutionary conservation, a popular feature in many other functional-impact predictors, has almost no contribution. Moreover, TF binding inferred from *in vitro* SELEX is as effective as that from *in vivo* ChIP-Seq. Second, we implemented GRAM integrating SELEX features and expression profiles. The program combines a universal regulatory score for a variant in a non-coding element with a modifier score reflecting the particular cell type. We benchmarked GRAM on a large-scale MPRA dataset in the GM12878 cell line, achieving a ROC score of ∼0.73; performance on the K562 cell line was similar. Finally, we evaluated the performance of GRAM on targeted regions using luciferase assays in MCF7 and K562 cell lines. We noted that changing the insertion position of the construct relative to the reporter gene gives very different results, highlighting the importance of carefully defining the functional target the model is predicting.

**Author Summary:** Noncoding variants lie outside of protein-coding regions, and are found to have disease associations. However, knowledge on the molecular effect of these non-coding variants in a cell-specific context is very limited. Also, different output between multiple experiment platforms may introduce extra complexity in analyzing the molecular function of these variants. We developed GRAM, a generalized model to predict molecular effect of non-coding variants in multiple cell types for different experimental platforms. We first selected the most informative cell-independent SELEX transcription factor binding score on the variant locus as features and then combine cell-specific gene expression profile to build a multi-step prediction model. GRAM has been successfully tested on both MPRA and Luciferase assay, and on three different cell lines: GM12878, K562 and MCF7, shows high performance.

## Introduction

Advances in next-generation sequencing (NGS) technologies have enabled high-throughput whole genome and exome sequencing [1], which has led to the identification and characterization of many disease-associated mutations [2] and the vast majority of common single nucleotide variants (SNVs) in the human population [3, 4]. Genome-wide association studies (GWAS) have found that these variants mostly lie outside of protein-coding regions [5], emphasizing the functional importance of non-coding regulatory elements in the human genome. This also defines an urgent need to develop high-throughput methods to sift through this deluge of sequencing data to quickly determine the functional relevance of each non-coding variant [6].

Evidence suggests that only a fraction of non-coding variants are functional, and the majority of functional variants show only modest effects[7]. Therefore, highly quantitative assays are needed to study large numbers of variants. Luciferase reporter assays are a common method to measure the regulatory effects of functional elements [8]. Comparing the difference of luciferase expression with and without a mutation can be used to estimate the experimental molecular effect of non-coding variants lying in a functional element. By using high-throughput microarray and NGS technology, the massively parallel reporter assay (MPRA) has extended the scales to the genome-wide level [9-14]. Recently, Tewhey and colleagues demonstrated the capability of MPRA to identify the causal variants that directly modulate gene expression [15]. This study identified 842 expression-modulating variants (emVARs) showing significantly differential expression modulation effects and provided a high-quality data source for computational modeling [15, 16].

There is an increasing need for computational methods to effectively predict the molecular effects of variants and improve our understanding of the underlying biology of these effects. Several approaches have been developed to address the problem of variant prioritization from different perspectives. Based on the target of predictions, these methods comprise two major categories: 1) disease-causing effect predictors (e.g., DeepSEA [17], GWAVA [18] and CADD [19]), which aim to prioritize causal disease variants and distinguish them from benign ones; and 2) fitness consequence prioritization (e.g., fitCons [20] and LINSIGHT [21]), which attempt to identify the variants based on evolutionary fitness. Other tools (e.g., Funseq2 [6]) do not belong to these categories because they integrate a comprehensive data context and unsupervised scoring system [6]. These computational methods are designed to predict and prioritize deleterious and disease-associated variants, but not specific molecular phenotypes of these variants (i.e., their effects on the activities of functional elements). Most importantly, none of the above tools take into account cell specificity in their models. One reason may be that some cell-specific features derived from chromatin immunoprecipitation sequencing (ChIP-Seq) data are only available in a few cell lines, which is a major hindrance to the generalization of a model.

In this study, we addressed the problem of molecular effect prediction of variants from a new perspective. Instead of predicting phenotypic consequences from genotypes, which is a common practice, we aimed to directly predict the expression-modulating effect of the variants from various sources of information. Our model incorporates selected transcription factor (TF) binding information from *in vitro* SELEX assays, representing the general binding potential of TFs on the variant’s location, and cell type-specific expression profiles, representing cellular contexts. Combining cell type-independent and-dependent features gives our model both flexibility and specificity. Evaluation results from MPRA and luciferase assay experiments show our model achieved high predictive performance and can be easily transferred to other cell types and assay platforms.

## Results

### Overall analysis flow

In this study, we first collected a dataset from Tewhey et al. [15] for estimation of expression modulation differences between wild type and mutants in the GM12878 cell line. This MPRA-generated dataset contains 3,222 SNVs filtered by logSkew value, which measures the log-fold change of the expression-modulating differences between wild-type and mutant alleles. Among them, 792 variants (named emVARs) had a significant expression-modulating effect compared with their respective wild-type allele, which indicates the molecular effect of the variant. Here, we treated emVARs and non-emVARs as a positive and negative dataset in our GRAM model.

As described in Fig. 1, our GRAM model is implemented with three steps: the first step predicts the universal regulatory consequences of the element with variant using SELEX TF binding score; the second step combines TF binding score with cell type-specific TF expression profiles to predict the cell type modifier score in a specific cellular context; the third step integrates outputs from the previous two phases to estimate the expression modulating effect in a cell-type-and assay-platform-specific context.

**Fig. 1:**
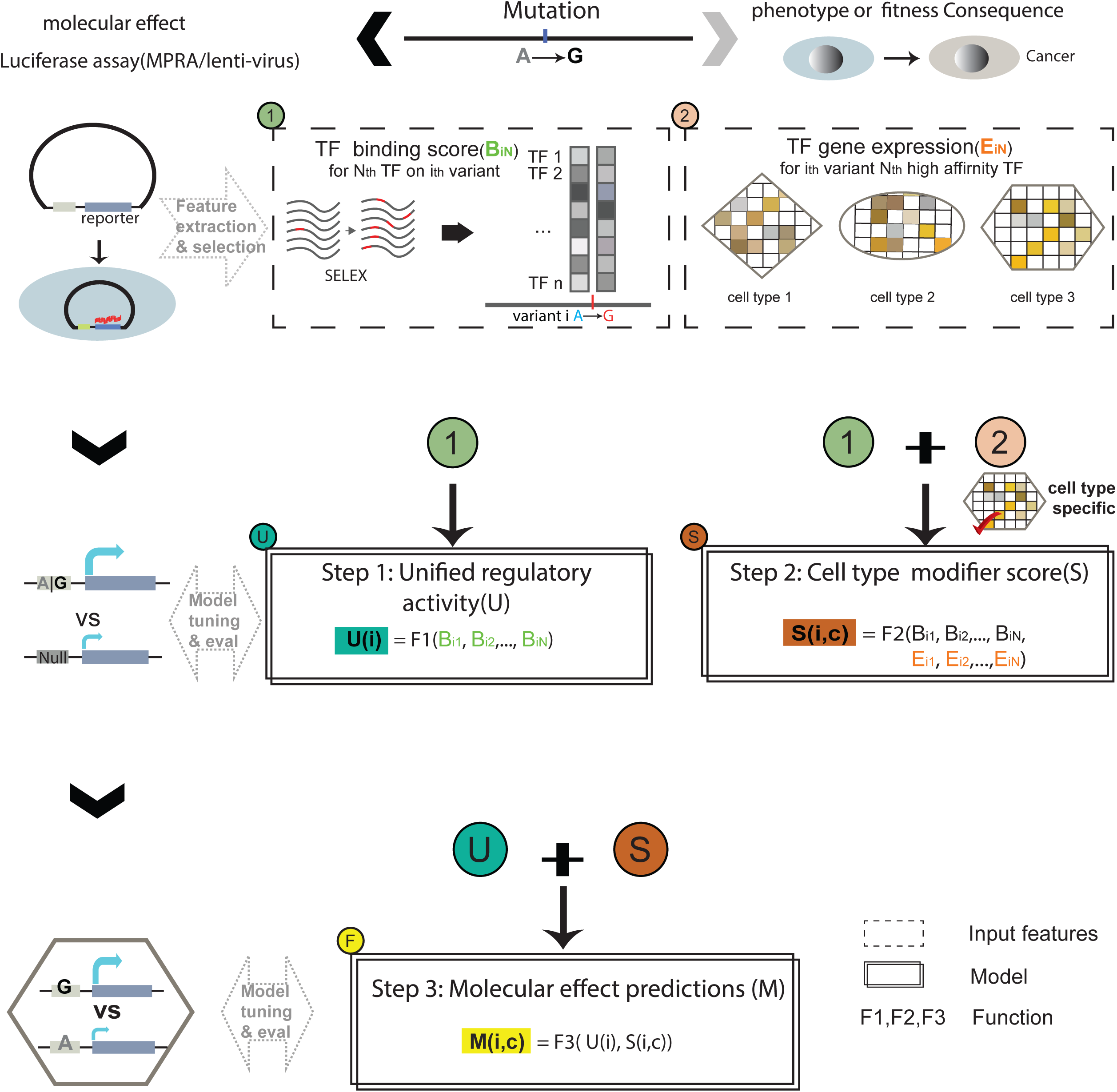
Overall flow of GRAM. The model predicts functional effects given the genotype with three steps: the first step predicts a universal regulatory activity using TF binding features; the second step predicts a cell type-specific modifier score using TF binding score and expression profiles; the final step integrates the results from the previous two steps to predict the expression-modulating effect of the variant.

### Exploring conservation and TF binding features

We first investigated the potential of evolutionary conservation and transcription binding features as predictors. Evolutionary conservation is associated with deleterious fitness consequence and is widely used in prioritization algorithms of non-coding variants, such as PhyloP [22] and PhastCons [23] score in LINSIGHT [21] and CADD [19], and GERP [22] score in Funseq2 [6]. We performed comparative analyses for these three conservation features across different datasets (Fig. 2a). We found that the PhastCons and PhyloP pattern of emVARs and non-emVARs are different from Human Gene Mutation Database (HGMD) [24] variants but similar to non-HGMD variants, which are thought to be benign. GERP scores show a similar pattern but have smaller variance in emVARs and non-emVARs compared to other datasets, with slightly larger values for emVARs. As we did not find differential patterns when comparing emVARs and non-emVARs, we further discovered that the correlation between logSkew and conservation scores was low with the explained variance very close to 0 for all three evolution features by linear regression. These results indicate that these conservation scores have little or no contribution to molecular effects on their own.

**Fig. 2:**
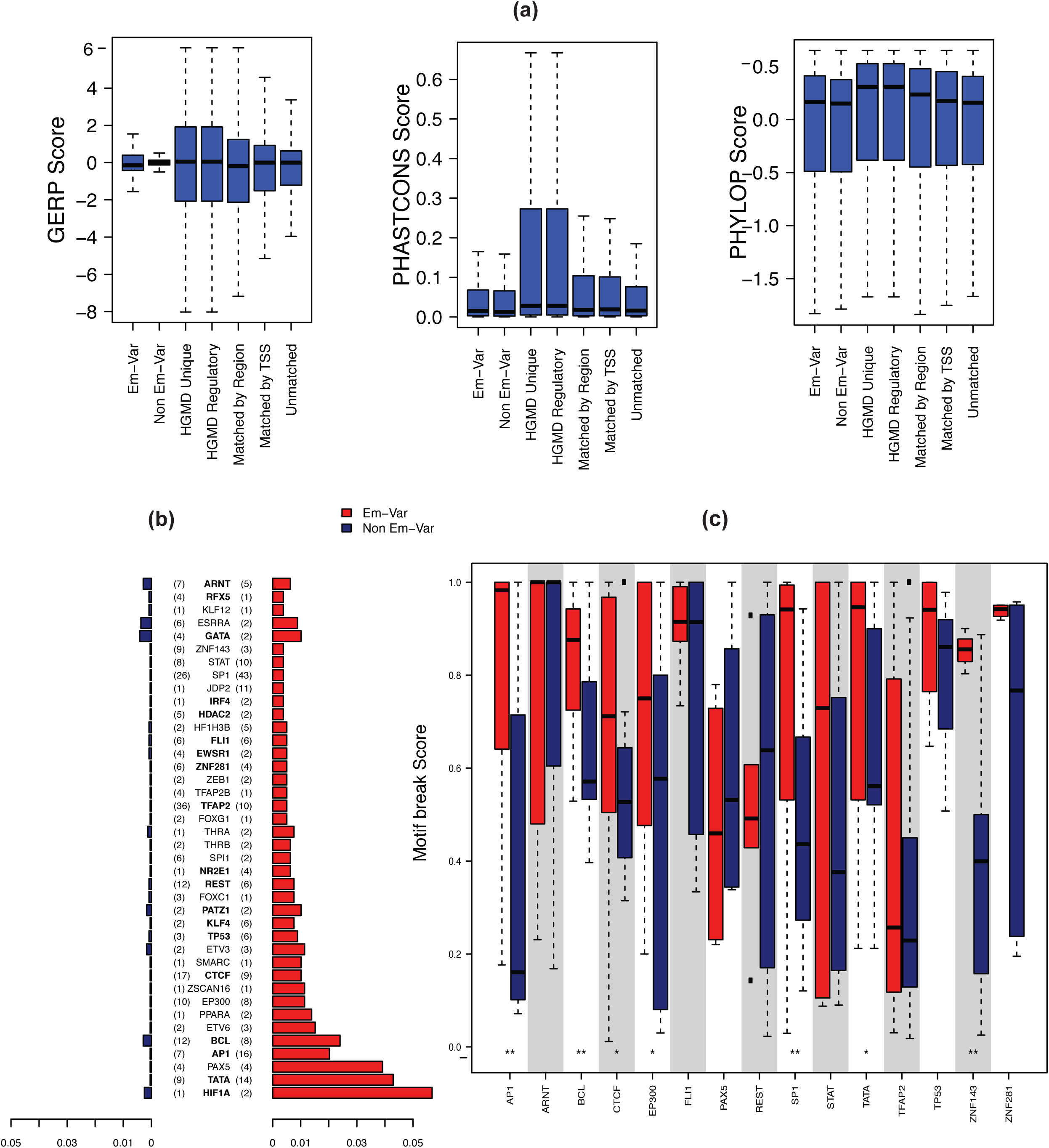
Preliminary selection of predictive features. (a) Distribution of conservation scores among different annotation categories. (b) Enrichment of TF binding peaks in emVAR and non-emVAR sets. x-axis is ratio of variants overlapping with the TF peaks over all variants in the same set. The TFs are sorted with p-values in hypergeometric distribution test in an decreasing order. The number in the bracket indicates the observed motif break event count. TFs with sufficient number observations are highlighted in bold. (c) Motif break scores in wild-type and mutant alleles for TFs with sufficient observed event count.

TF binding can link the molecular effect of non-coding variants to a cascade of a regulatory network, which is thought to be an important contributing factor to the variants’ regulatory effects [17, 19, 25, 26]. Tewhey et al. found that the logSkew value positively associates with TF binding scores. To thoroughly evaluate the effect of TF binding, we tested TF binding peaks overlapping with the SNVs and TF motif break events in the Tewhey dataset. We annotated and analyzed the emVAR and non-emVAR variant sets with Funseq2 [6], and found that the emVAR set had more TF binding events compared with the non-emVAR set (Fig. 2b). In addition to TF binding enrichment, we examined the motif break scores for these TFs. After removing TFs with insufficient observations, the differences between the distribution of motif-break score for mutant and wild-type genotypes in emVARs are larger than those in the non-emVAR dataset (Fig. 2c). According to this analysis, the emVAR set tends to have not only more TF binding events, but also larger binding alterations compared with the non-emVAR set. Thus, TF binding shows high association with the expression-modulating effects of the variants.

### Model-based feature selection

We generated TF binding features for potential training features using 515 Deepbind models inferred from both ChIP-Seq [27] and *in vitro* SELEX assays [28]. With a comprehensive feature selection framework to select impactful TF binding features, we prioritized these features across models with LASSO stability selection [29] and Random Forest (shown in Fig. 3a). The 20 most important features (out of 515) with respect to mean importance across all methods is shown in decreasing order in Fig. 3a. Both ChIP-Seq and SELEX Deepbind features showed high importance, with the top two being GM12878 ChIP-Seq features (SP1 and BCL3), which are cell line specific, followed by SELEX features starting with ETP63. The top-ranked impactful TFs tend to have more TF-TF interactions than the bottom-ranked TFs, indicating that the importance of a TF reflects its role in the TF-TF cascade regulatory network (Fig. 3b).

**Fig. 3:**
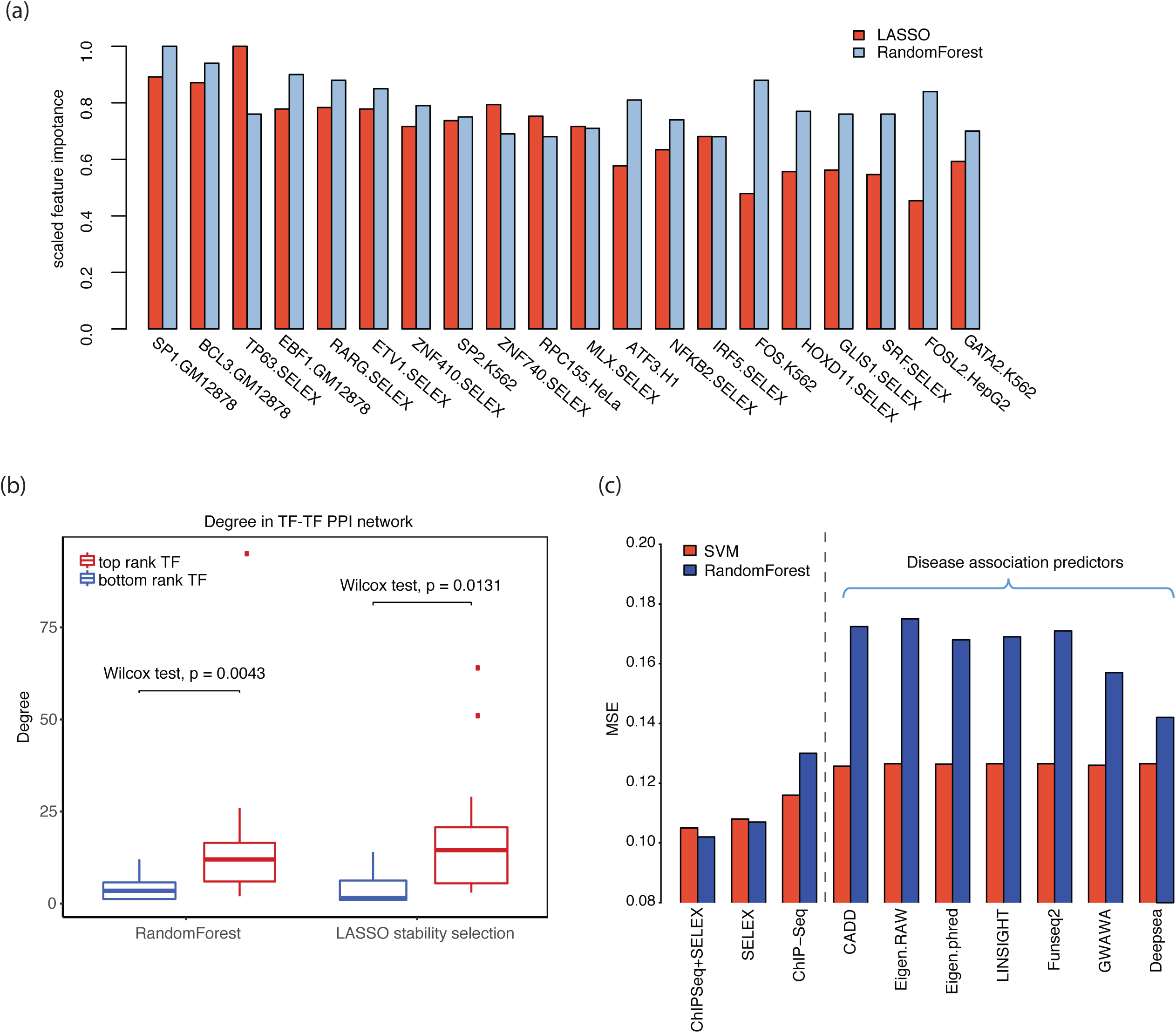
Model based feature selection. (a) Importance of the top-ranked features for SELEX-and ChIP-Seq-derived models. The features are sorted according to the mean of LASSO stability selection and random forest importance scores. (b) Regulatory network degree of relevant TFs for the top-ranked and bottom-ranked TFs in LASSO stability selection and random forest. (c) Comparison of the performance of different feature sets, including cell-line specific ChIP-Seq TF binding scores and SELEX TF binding scores, as well as features defined from previous disease-association prediction tools.

Interestingly, many SELEX features, though not cell type-dependent, achieved similar predictive power as cell type-specific ChIP-Seq features. We compared the predictive performances of cell type-dependent ChIP-Seq features, cell type-independent SELEX features, and combination of both feature sets using LASSO regressor, support vector machine (SVM) regressor and Random Forest. Incorporating ChIP-Seq-derived features did not boost the accuracy significantly for any of the three models (Fig. 3c and S1 Table). As the availability of ChIP-Seq is restricted to a few cell lines, we instead used SELEX features to build a generalized model across different cell types.

We then used the features generated from disease-association prediction tools (CADD [30], Funseq2 [31], DeepSEA [32], GWAVA [33], LINSIGHT [34], and Eigen [35]) to predict the same molecular effect target. As shown in Fig. 3c, this analysis indicated that the prediction of disease-associated variants is not equivalent to that of expression modulating variants.

### Building a generalized model by multi-step learning

Using the TF binding features from DeepBind models and the MPRA dataset from Tewhey et al. [15], we implemented our multi-step GRAM model. In the first step, we predicted the universal regulatory activity of an element with or without a variant. The 10-fold cross validation demonstrated exemplary performance of the model with an area under the receiver operating characteristic curve (AUROC) of 0.938 and an area under the precision-recall curve (AUPRC) of 0.924 (Fig. 4a and S1 Fig).

**Fig. 4:**
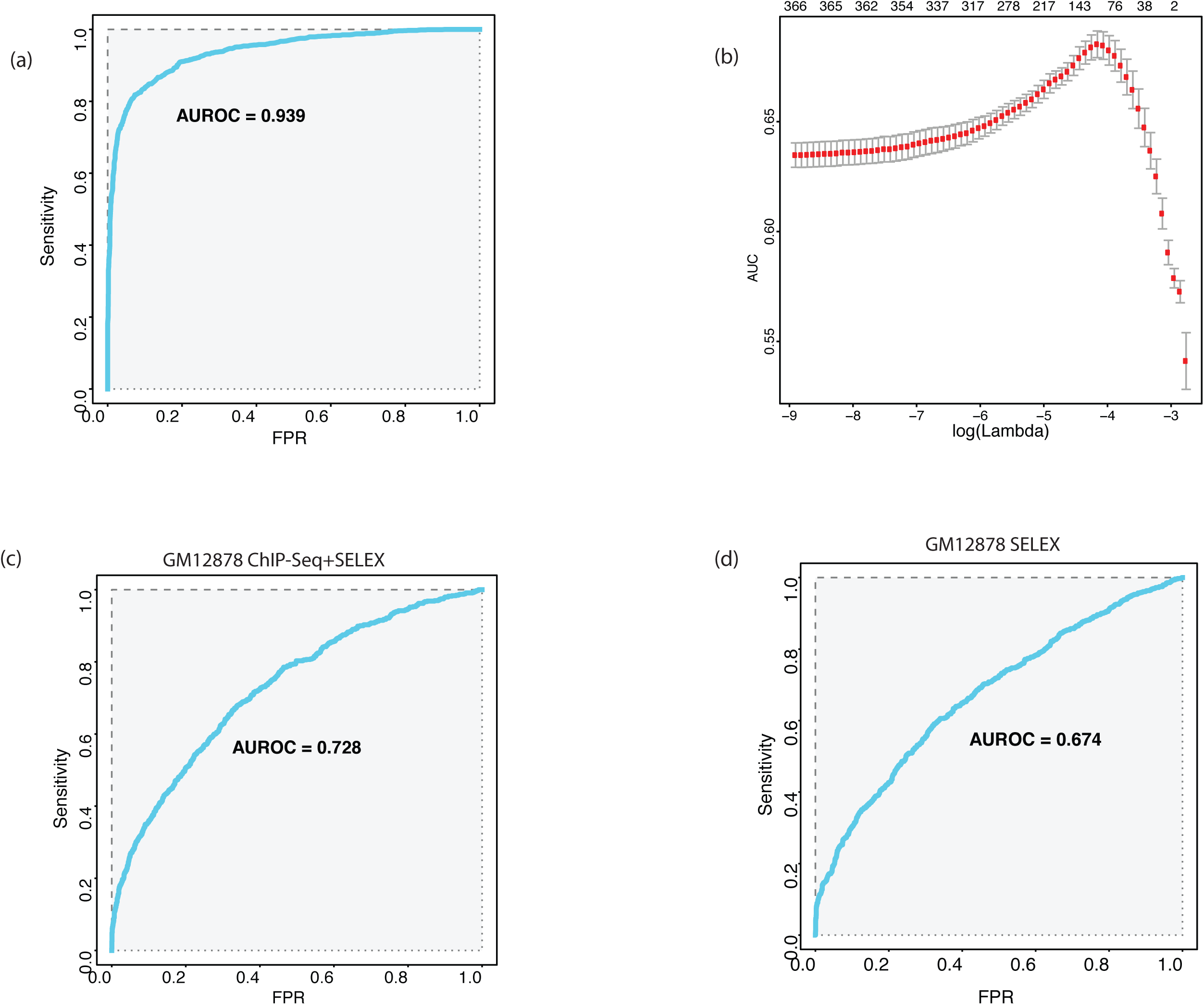
Performance of the GRAM multi-step model. (a) ROC curve for regulatory activity prediction. (b) The prediction of cell specificity using TF expression profiles. (c) (d) ROC for model with/without tissue-specific ChIP-Seq Deepbind features.

In the second step, we calculated a cell-type modifier score as an indicator of the experimental assay’s cell-specific nature (see Methods for details). Briefly, we defined the prediction target using a top and bottom quantile of Vodds (S3 Fig). Vodds is the standard deviation of log odds for each variant’s read count in MPRA and it shows cell line-specific patterns (S2 Fig). We found that variants with higher Vodds tend to include more non-emVARs (Chi-square test p-value: 0.0002). Hence, the cell type modifier score can be used to adjust the universal regulatory effect to a cell type-specific context.

Gene expression profile, especially TF expression, are more generally available and can represent the cellular environment. We incorporated TF gene expression and TF binding scores as features to predict the cell type modifier target, and got an AUROC of 0.65 and 0.8, respectively, using LASSO 10-fold cross-validation (S4 Fig and Fig. 4b).

The final step of our GRAM model is to predict the molecular effect of a variant that can significantly modulate reporter gene expression. To do this, we fed the output from the first and second step into a LASSO model, with the emVAR and non-emVAR labels as targets. We found that the AUROC of 10-fold cross-validation for the optimal model is 0.728 (Fig. 4c) and the AUPRC was 0.505, which are higher than the state-of-the-art method (KSM) using the same dataset (AUROC: 0.684, AUPRC: 0.478) [36].

We also tried to build a generalized model by removing cell type-specific ChIP-Seq features. We repeated step one and two on the same dataset using GRAM cell-independent features from the SELEX assay, which achieved comparable performance with AUROC = 0.674 (Fig. 4d) and AUPRC = 0.452.

### Validating the GRAM model using experimental assays

We next evaluated performance of the model on different cell types and assay platforms. We use the generalized model trained on GM12878 cells and tested it on another MPRA dataset in K562 cells [10], which includes 2,400 elements in 149bps with a variant centered on the inserted fragment (Fig. 5a-b). The AUROC for Step 1 universal regulatory activity was as high as 0.68 when we set the q-value cutoff to 10^−9^; the molecular effect prediction was also over 0.8 if we used a more stringent q-value cutoff (10^−5^).

**Fig. 5:**
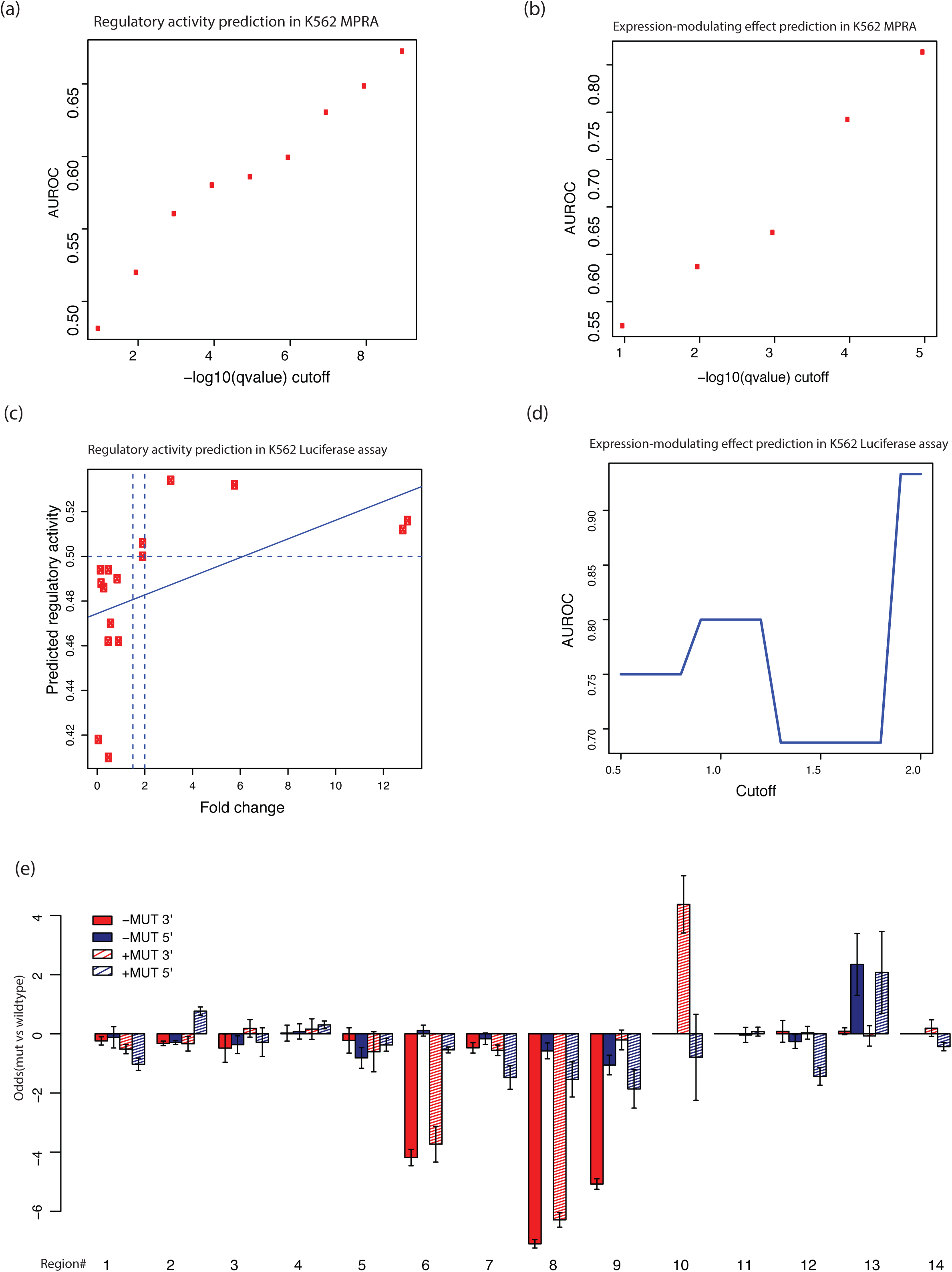
Experimental validation. (a) Regulatory activity prediction results on an independent K562 MPRA dataset (b) Expression-modulating effect prediction results on an independent K562 MPRA dataset. (c) Regulatory activity prediction for luciferase assay in K562. The x-axis represents fold change from the experiment. The vertical dot lines represent the cut off (1.5, or 2) to determine positive (enhancer) and negative, and the horizontal dot line is the predicted probability cutoff (0.5). (d) The AUROC value versus the different absolute log2 odds cutoff [0.5, 2.0]; (e) Experimental results (in odds ratio) for luciferase assay in K562. The 5’ terminal and 3’ terminal insertions are compared.

Other than measuring read counts as in MPRA, some other assays, such as luciferase and GFP reporter assays, measure luminescence and fluorescence readouts instead. [37, 38]. To evaluate how our model, trained with MPRA data, can be transferred to these assay platforms we tested its performance on luciferase assay results of eight potential regulatory elements with mutations from the MCF7 cell line [39]. For regulatory activity, the predicted probability of being an active regulator was positively correlated with luciferase assay fold change between the inserted element and background control. The results show perfectly prediction (AUROC=1) using fold change cutoffs from 1.2 to 2 (Fig. 5c). For the prediction of expression-modulating effects, we defined the significant changes between mutant and wild type by using an absolute log2(odds ratio) cutoff. The predicted probability also showed a positive correlation with absolute log2 fold change (S5 Fig). The AUROC value ranged from 0.7 to 0.9 given the absolute log2 cutoff from 0.5 to 1.5 (Fig. 5d). This indicates that our model performs very well on luciferase assay or MPRA dataset from different cell lines, though these assays use different measurements.

In MPRA, the element is inserted upstream (5’-terminal) of the reporter gene, but for some assays, such as STARR-Seq, the element is inserted downstream (3’-terminal). Therefore, we further tested the effect of insertion location of an element in luciferase reporters in K562 cells using 14 randomly selected elements with potential regulatory activity. As shown in Fig. 5e, the 5’ terminal log odds were similar to the 3’ terminal odds for region 3, 4, 5, and 13, but showed significant differences for region 6, 8, 9, 10, and 14. The prediction of GRAM for the 5’ terminal was much better than that for the 3’-terminal insertions; the AUROC was 0.25 higher for universal regulatory activity and 0.32 higher for the expression-modulating effect prediction, indicating different mechanisms for the two ends. Therefore, GRAM is optimal for 5’ terminal assays.

## Discussion

There has been an increasing number of computational methods can prioritize non-coding variants. Also, accumulating high-throughput whole-genome sequencing data have become the primary source for identifying disease-associated variants. However, we still lack a tool that can estimate the expression-modulating effect of a variant in a cell-specific manner. In this study, we developed a multi-step generalized model called GRAM that can specifically predict the cell-type specific expression-modulating effect of a non-coding variant in the context of a particular experimental assay.

In this paper, we aimed to precisely define the expression-modulating effect as a function of the response variables extracted from genomic data. Unlike other variant impact prediction methods, we did not include evolutionary features in our model because it had very limited impact on the performance. We selected a variety of TF binding features that could be useful for predicting variant effects. We used direct measurements from TF binding and a straightforward LASSO regression to assess the importance of each feature. We found that *in vitro* SELEX TF features (aka non-cell-specific features) can achieve high predictive performance, which was further validated by SVM and Random Forest.

The three-step GRAM model predicts the expression-modulating effects of variants by integrating two intermediate predictive targets: universal regulatory activity and cell type modifier score. The universal regulatory activity reflects the regulatory effect of an element with/without a mutation in a vector-based assay without cell type-specific chromatin contexts and epigenomics information. Since cell-specific information cannot be ignored for predicting the variant’s effects, we further adjusted the universal regulatory effect with the cell type modifier score in the final step of the prediction model, resulting in much better performance by the model. We note that our framework can be further converted to a Bayesian hierarchy model with the intermediate targets as hidden variables.

GRAM performed well with targeted validations on MPRA and luciferase assay platforms, even across different cell types. In addition to target validations, we could explore in great detail the sensitivity of these methods and the impact of vector construct. The insertion position of the element affected the outcome of the assay, which may correspond to different types of regulatory elements. Because our model is trained on 5’-terminal insertion data, the prediction is consistent with outcomes from the same position, but not for 3’-terminal assay results. This indicates different mechanisms for two insertion positions: an element inserted upstream of a reporter gene may detect either the promoter or enhancer activity of the element. On the other hand, if it is inserted downstream of the gene’s transcriptional start site or the 3’ terminal, it may specifically target to the enhancer activity. However, large-scale experimental validation is required to further elucidate the underlying mechanisms.

The GRAM model will be a useful tool for elucidating the underlying patterns of variants that modulate expression in a cell-type context. By leveraging the accumulating data generated from multiple cell lines, future studies can be performed in-depth investigation using GRAM. We will keep abreast with the growing availability of comprehensive datasets and further expand our analysis.

## Methods

### Dataset

We downloaded the dataset from R. Tewhey et al.’s paper [15]. From about 79K tested elements, we only kept variants for which either wild-type or mutant elements show regulatory activity. This reduced the set to 3,222 SNVs in the GM12878 cell line. Each SNV was extended in both directions by 74bp, in total 149bp. We used another dataset from Ulirsch 2016 [10], which included 2,756 variants tested in the K562 cell line.

The protein-protein interaction network used in our downstream analysis was constructed by merging all interaction pairs identified by BioGrid [40], STRING [41] and InBio Map [42].

### Feature extraction

GERP features were extracted using the Funseq2 annotation pipeline, which searches the region of elements over the whole genome GERP score file to get an average score. We downloaded phyloP [22] and Phastcons [23] scores from the UCSC genome browser data portal (http://hgdownload-test.cse.ucsc.edu/goldenPath/hg19/).

We performed motif enrichment analysis using a hypergeometric test. For the motif break and gain score comparison, we removed the TFs that covered less than two variants for either emVARs or non-emVARs from the top 40 TFs. Then, we performed a Wilcoxon test for the motif break score.

Motif break and motif gain scores were calculated using Funseq2. We also calculated the motif score using Deepbind [26] with both the SELEX and ChIP-Seq motif model. The SELEX motif model is based on an *in vitro* binding assay: systematic evolution of ligands by exponential enrichment. However, ChIP-Seq models are inferred using sequences from the TF binding site from different cell lines. A total of 515 motif models were calculated (table s1: tbls1.deepbind.list.txt).

### Model-based feature selection

To examine the importance of features, we compared different metrics, which included LASSO stability selection [29] and Random Forest regression. The feature importance for each selection method was scaled to [0, 1]; we took the mean of all the selection methods to represent the overall ranking.

We compared our models’ mean standard error (MSE) with CADD, Eigen, LINSIGHT, Funseq2, GAWVA, and DeepSea. Features from the above tools were collected and tested using both SVR and Random Forest regression, which considered all Deepbind features and SELEX-based and ChIP-Seq-based features, respectively. For the other variant prioritization tools, we took the output of these methods, and then used the same SVR and Random Forest to train and predict the logSkew value.

### GRAM – multi-step generalized model

Pseudocode:

i: variant id

j: TF id

M: the total number of variants

N: the total number of TF

*B*_*ij*_: TF j binding score on *i* variant

*E*_*ij*_: Expression of jth high-affinity TF on *i* variant

c: cell type

U(i), S(i,c), F(i,c): model for step 1-3

Step1: simple Universal score to be a regulatory element using randomForest classifier, *U*(*i*) ∈ [0,1]

*U*(*i*) = *F*1 (*B*_*i*1._, *B*_*i*2_, …, *B*_*iN*_)

Step2: cell type modifier score, *S*(*i, c*) ∈ [0,1]

*S*(*i, c*) = *F*2(*B*_*i*1._, *B*_*i*2_, …, *B*_*iN*_, *E*_*i*1._, *E*_*i*2_, …, *E*_*iN*_) = *logit*(*b*_*i*1._*B*_*i*1._ + *b*_*i*2_ *B*_*i*2_ + *…* + *B*_*iN*_*T*_*iN*_ + *b*_i′1_*E*_i1._ + *b*_*i*′2_*E*_*i*2_ + *…* + *b*_*i*′*N*_*E*_i′*N*_+ *b*_*ic*_)

Objective function: 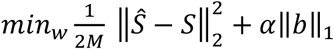

Step2: molecular effect score, *S*(*i, c*) ∈ [0,1]

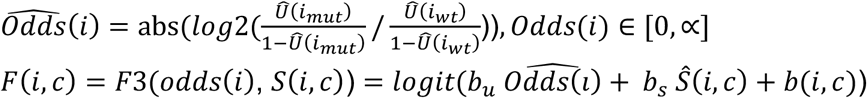

Objective function: 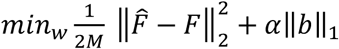

We defined the emVARs as positive and non-emVARs as negative classes following [15], where ‘expression modulating’ means having a molecular effect that significantly increases or decreases regulatory activities. In total, we used 3,222 data records, including 799 positive and 2,423 negative.

We built a three-step GRAM model. Model 1 predicts the universal element regulatory activity *U* for wild type and mutant. The ground-truth of regulatory activity is determined by experimental assay platforms, like luciferase assay or MPRA. An element inserted into plasmid with or without a mutation is defined as a regulatory element if the fold change between the element and the control is larger than a statistically significant cutoff. For example, for the MPRA study, a statistical test based on DESeq2 indicates whether it is significantly changed; for the luciferase assay, we considered a testing element that has a fold change greater than 1.5 or 2 compared to control (eGFP) as a regulatory element. The response variable is the TF binding score changes from wild type to mutant, which is estimated by Deepbind. A Random Forest classifier was trained to predict the universal regulatory activity. The predicted log odds of probability between the wild type and mutant was calculated by 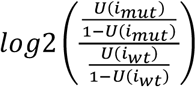

Model 2 predicts cell type modifier scores using gene expression and binding scores of TF. The cell type modifier score is defined according to the cell specificity of the experimental assay. Given the reads count of MPRA for wild type, mutant, and their null control, it forms a 2×2 categorical table. The standard deviation of log(odds) of the categorical table (n1,n2,n3,n4 for the average reads count) is calculated as 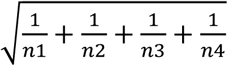 By comparing principal component loading of the Vodds from three cell lines: GM12878, GM19239, and HepG2, we found that the two GM cell lines are closer to each other than to HepG2 (Fig. S2), which indicates that Vodds contains cell type information. The underlying biology may reflect the cell specificity of the experiment, such as the success rate of transfection. We then further compared two groups of variants with the top quantile and bottom quantile of Vodds in GM12878, and found that there were more non-emVAR variants in the top quantile group, which indicates the Vodds are also associated with the molecular effects of the variants. Then, we defined a cell type modifier target using the top and bottom quantile variants.

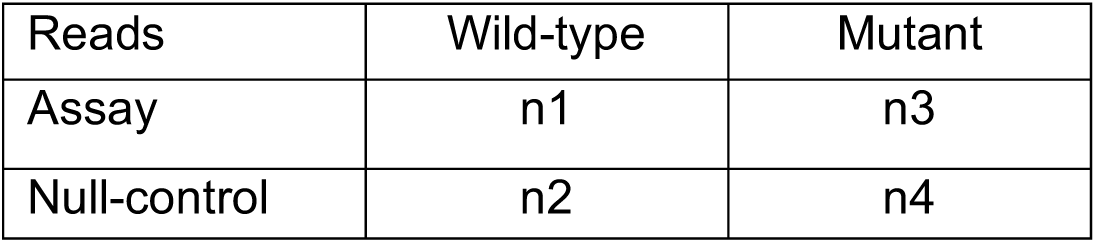

| Reads | Wild-type | Mutant |
| --- | --- | --- |
| Assay | $n_1$ | $n_3$ |
| Null-control | $n_2$ | $n_4$ |

The TF expression profile was used to predict cell type index modifier class. For each mutation region, we adjusted the expression based on the TF binding score. Given 258 TFs with a Deepbind SELEX model score S for 3,222 SNVs, the TF expression matrix for each variant was adjusted and shuffled using the rank of SELEX TF binding scores among all the SNVs. Then, we used the TF binding score and gene expression to predict the cell type modifier class.

The final model predicts the molecular effect of a variant using the estimated universal odds ratio and cell type modifier from TF binding and expression as predictors. A simple LASSO was used to build the model.

### Experiment validation on MCF7 cells

Each regulatory region (both wild and mutant types) was separately synthesized. Enhancer regions were designed in such a fashion where based on the candidate SNV site, 250bp upstream and 250bp downstream was included for each enhancer region. These regions were then cloned into the pGL4.23[luc2/minP] vector (Promega, Cat# E841A). Each candidate region was placed upstream of the minP promoter to determine the effect of each putative enhancer region on luciferase expression. 100ng of each candidate construct and 100ng of Nano-luc control was co-transfected into MCF-7 cells (5,000 cells per well in DMEM media containing 10% FBS and 1% Penicillin-Streptomycin antibiotic) using the Lipofectamine 3000 reagent (Thermo Fisher, Cat# L3000001) according to manufacturer’s instructions. Cells were incubated for 48 hrs before reading the luciferase signal using Promega Nano-Glo luciferase kit (Promega, Cat# N1521) according to manufacturer’s instructions.

### Model validation using MPRA data from K562 cells

#### Enhancer Selection

Based on the enhancer prediction and histone mark signaling overlap, we randomly selected 14 putative regulatory elements, and then randomly picked one or two mutations based on Funseq2 whole genome scores (http://funseq3.gersteinlab.org). Next, we used a web tool to design site-directed mutagenesis primers to introduce the target SNVs into the 14 elements. Two SNVs were introduced into each element, with only one predicted to result in a significant change in enhancer activity.

#### Reporter Generation

Elements were amplified via PCR from human genomic DNA (Promega) with Platinum SuperFi polymerase (Invitrogen) and primers containing attB1 and attB2 sequences (see S2 Table). Elements were then cloned into pDONR223 using Gateway BP clonase and transformed into *E. coli* cells. Four colonies for each element were picked and sequenced via Sanger sequencing using the RV3 primer. One clone for each element with the correct sequence was then cloned into pDEST-hSCP1-luc or pGL4-Gateway-SCP1 using Gateway LR clonase, and luciferase reporters containing the elements were then transfected into K562 cells. pGL4-Gateway-SCP1 was a gift from Alexander Stark (Addgene plasmid # 71510) [44]. To construct a positive control for the enhancer activity assays, we cloned the widely used Rous sarcoma virus promoter that has been implied to possess enhancer activities.

#### Mutagenesis

The wild-type templates for site-directed mutagenesis were sequence-verified entry clones containing putative regulatory elements. The mutagenesis primers containing the pre-designed mutations were designed with a web tool (http://primer.yulab.org/). The mutagenesis reactions were carried out following the Clone-seq pipeline [43]. Each mutagenesis reaction contained a wild-type template and its corresponding mutagenesis primers. The products of the mutagenesis reaction were DpnI-digested and transformed into TOP10 chemically competent cells (Invitrogen). The transformants were spread on LB-spectinomycin agar plates and incubated at 37°C overnight. Single colonies yielded from the mutagenesis were picked, propagated, and sequence-verified before they were used in downstream experiments.

#### Cell Lines

K562 cells were a gift from the Melnick lab (Weill Cornell Medicine). Cells were cultured in Iscove’s Modified Dulbecco’s Medium (Gibco) supplemented with 10% fetal bovine serum and 1% Pen-Strep at 37°C with 5% CO2.

#### Luciferase Assay

K562 cells were transfected with 200 ng of the above reporters and 20 ng of Renilla luciferase (pRL-CMV, Promega) in triplicate in 96-well plates with Lipofectamine 3000 (Invitrogen). At 48 hours post-transfection, luciferase activity was assayed with the Dual-Glo Luciferase Assay System (Promega).

## Supporting Information

**S1 table.**
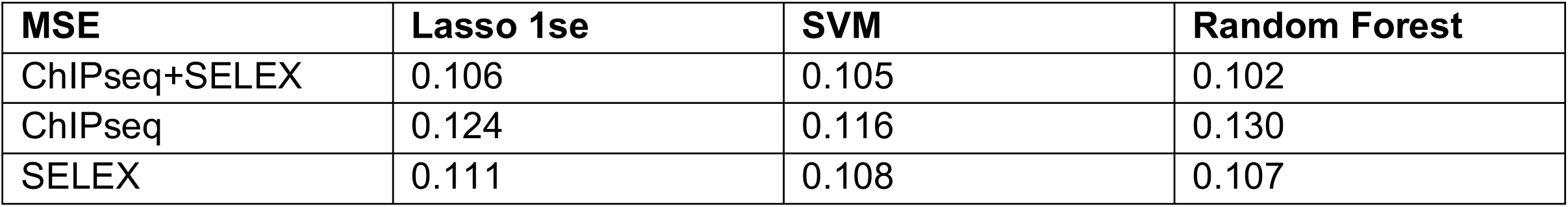
Predictive performance of different feature sets, including cell-line specific ChIP-Seq TF binding scores and SELEX TF binding scores, using Lasso, SVM and Random Forest

**S2 table.**
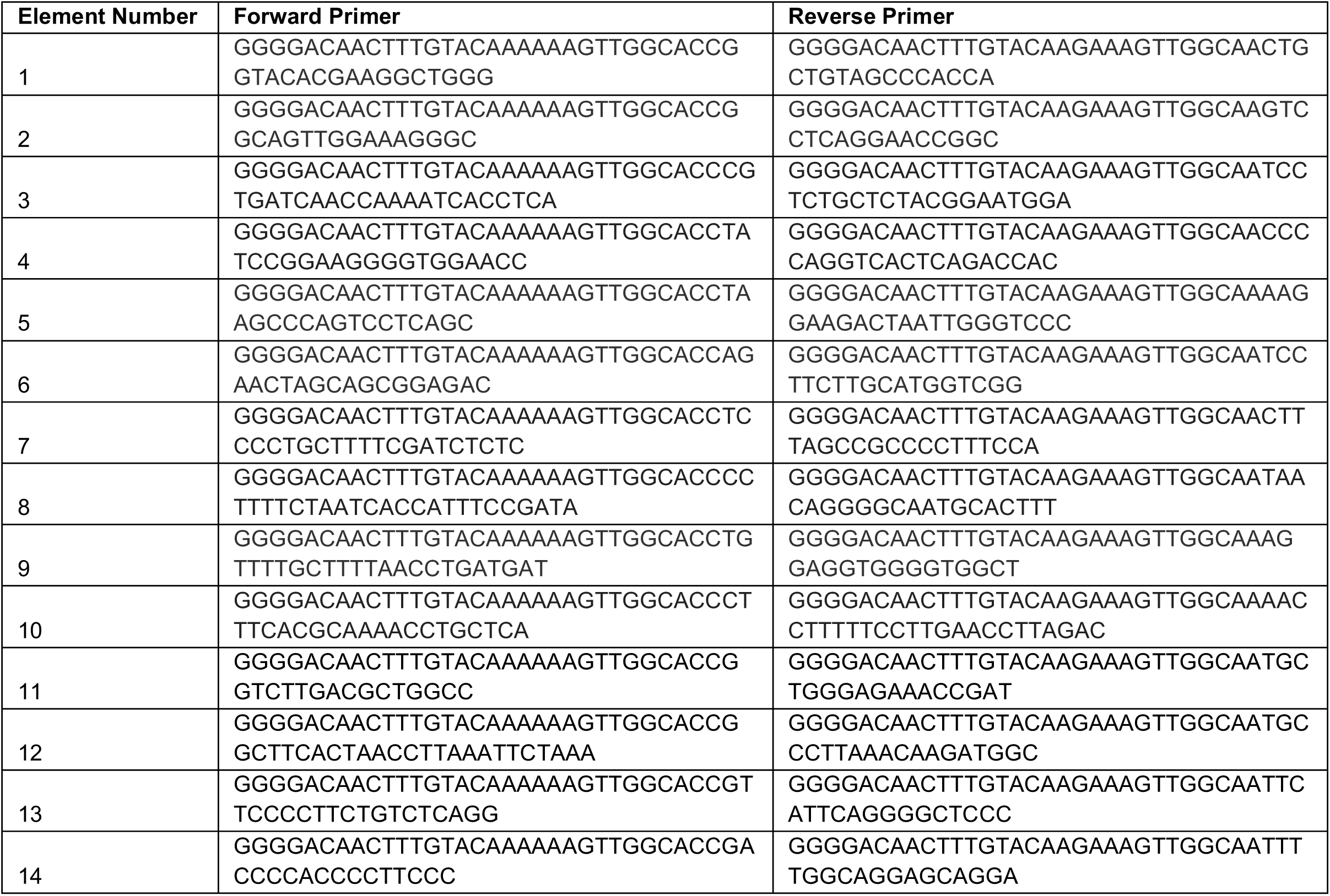
Primers for 14 regions cloning in K562

**S1 Fig.**
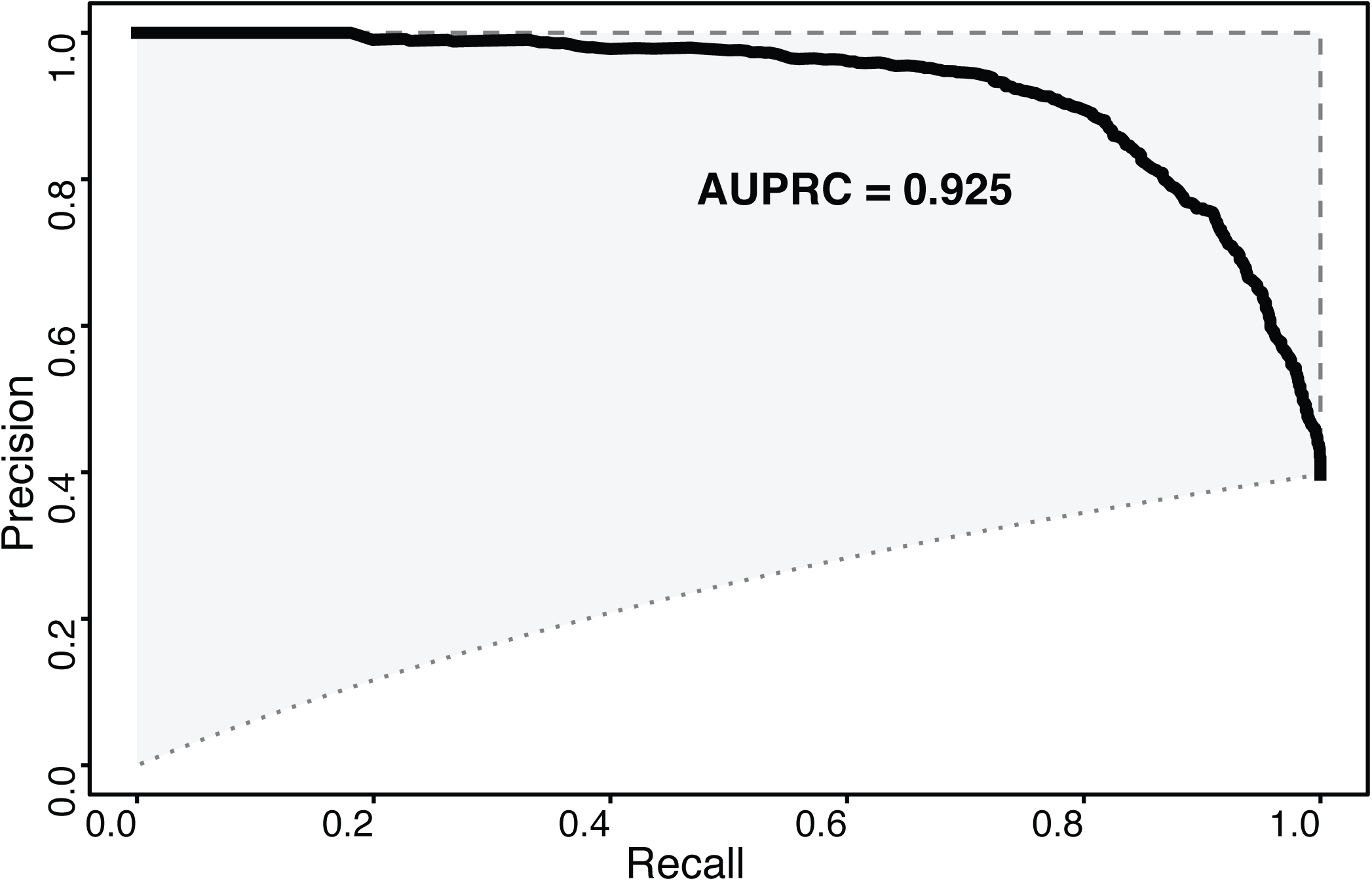
PRC curve for regulatory activity prediction

**S2 Fig.**
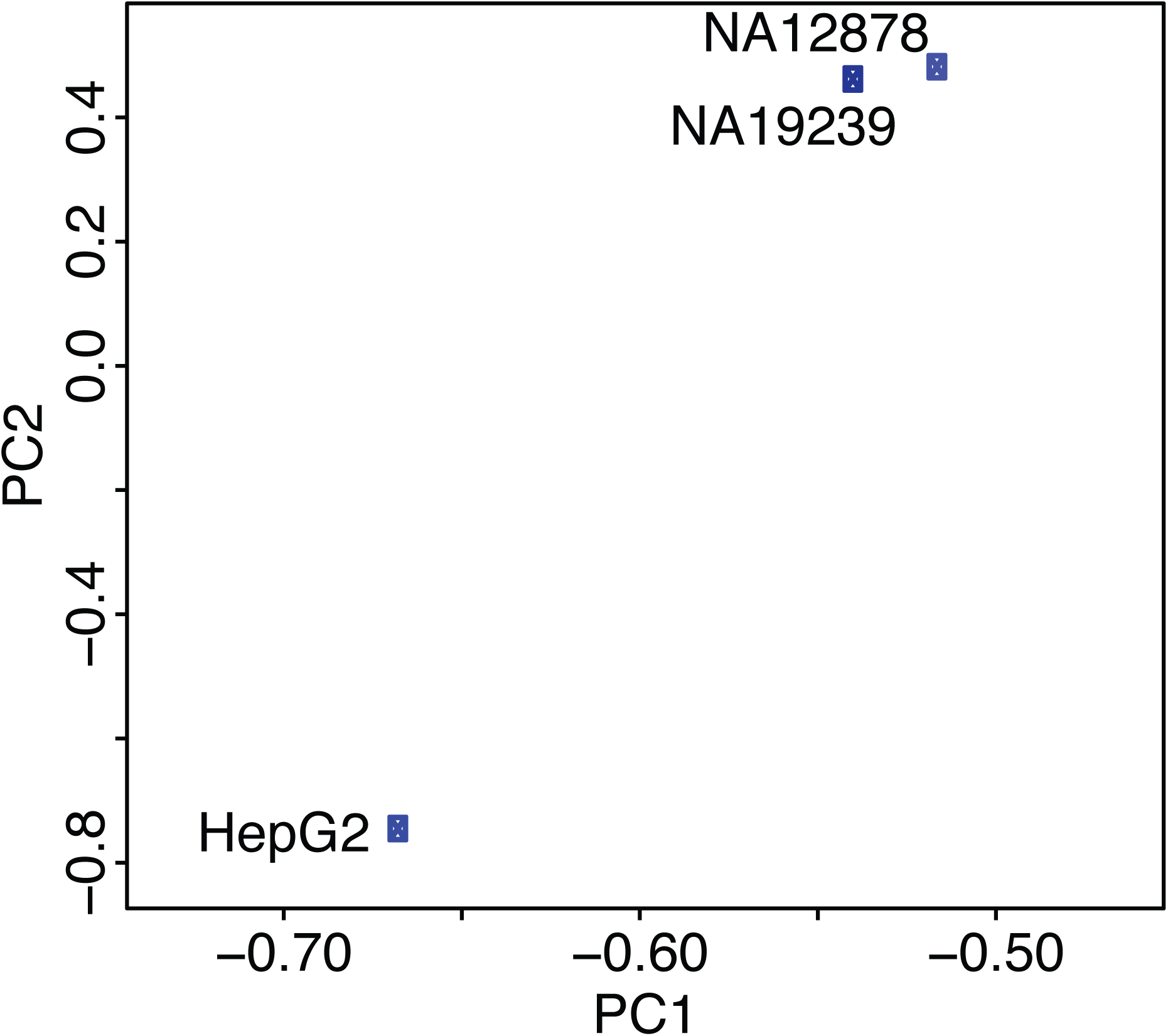
Principal component analysis using Vodds for three cell lines: GM12878, GM19239 and HepG2

**S3 Fig.**
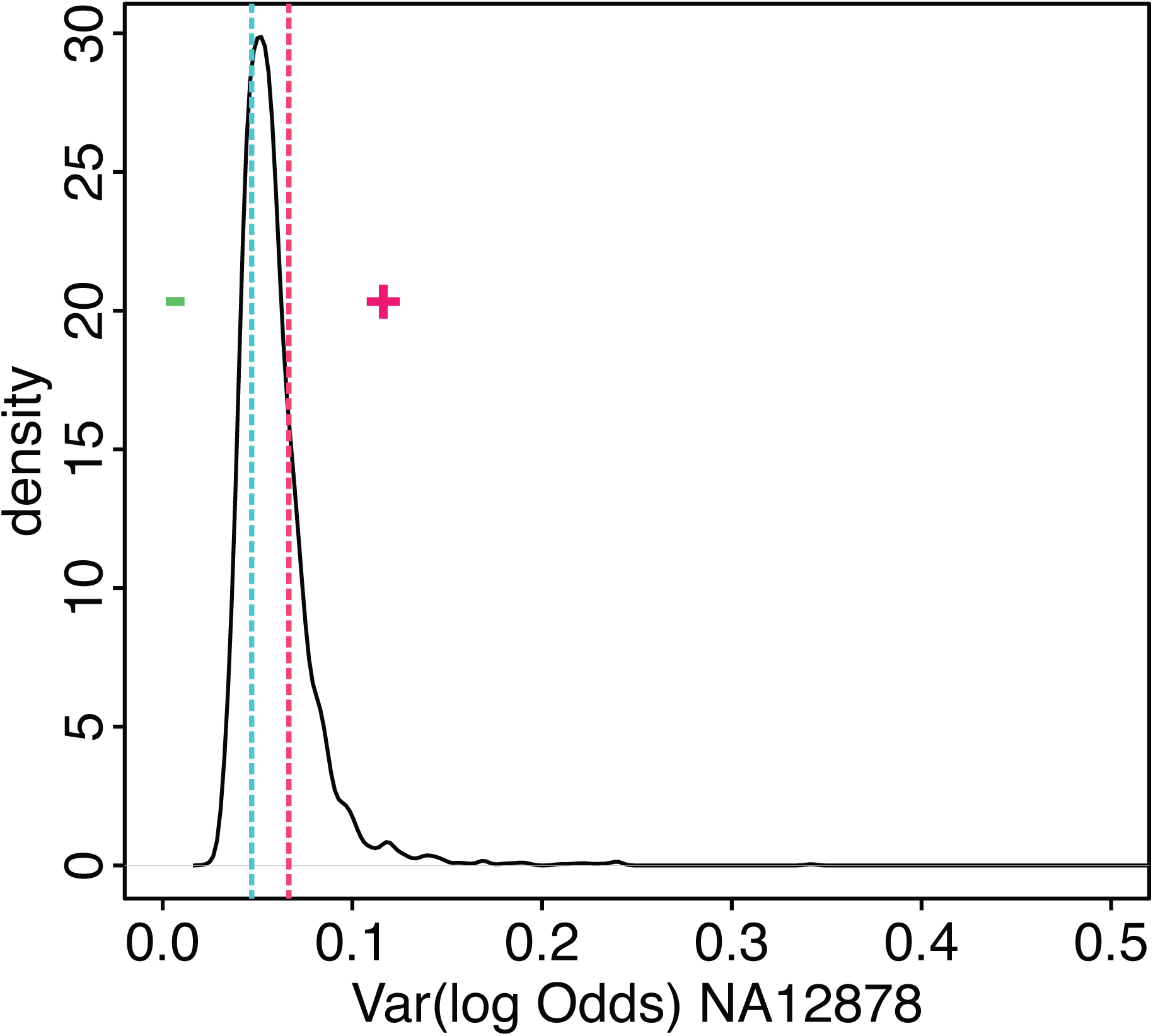
Distribution of Vodds score for GM12878. The high and low variable cell specificity class are defined by the top and bottom quantile.

**S4 Fig.**
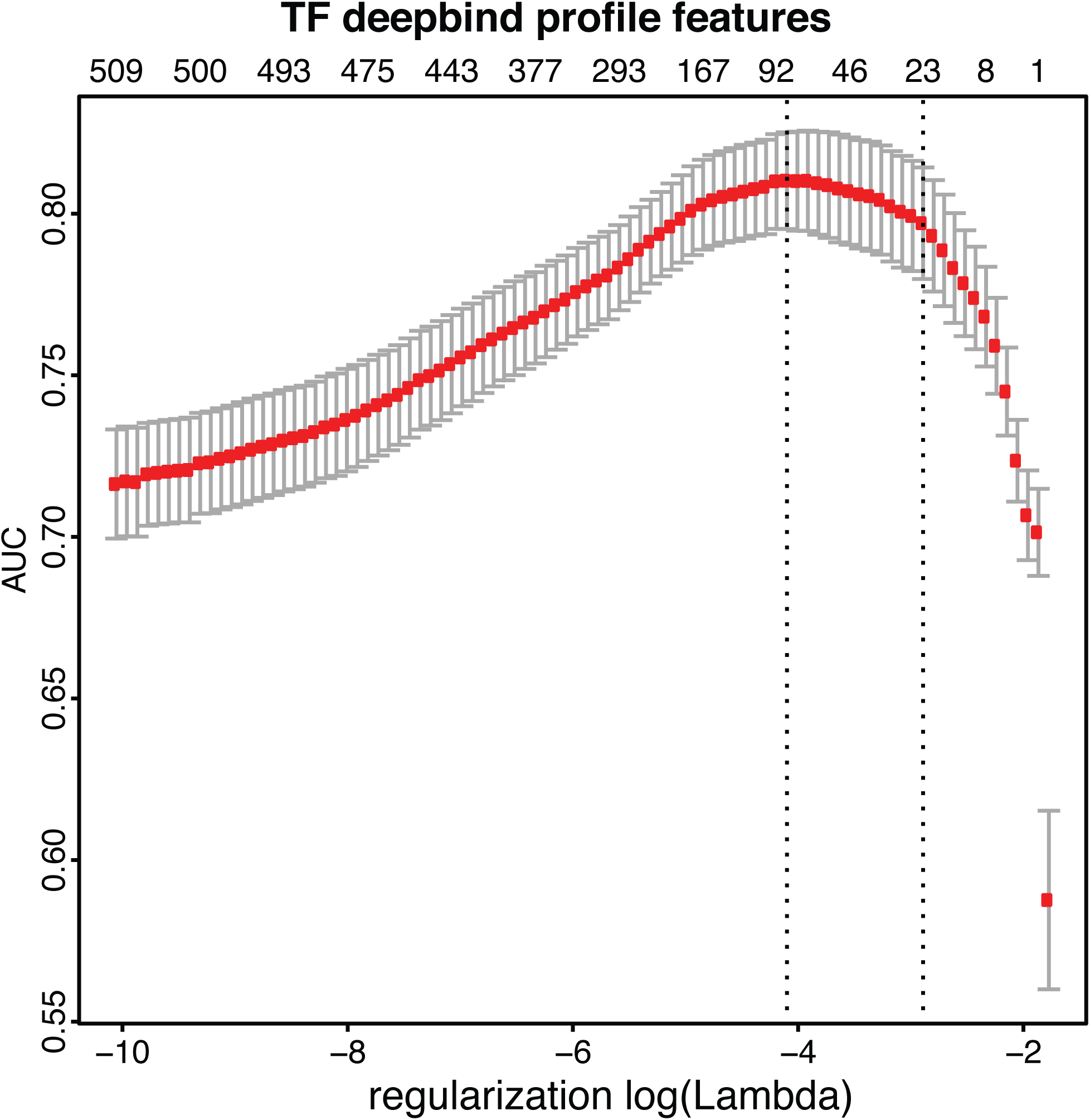
The prediction of cell type modifier score using TF binding profiles.

**S5 Fig.**
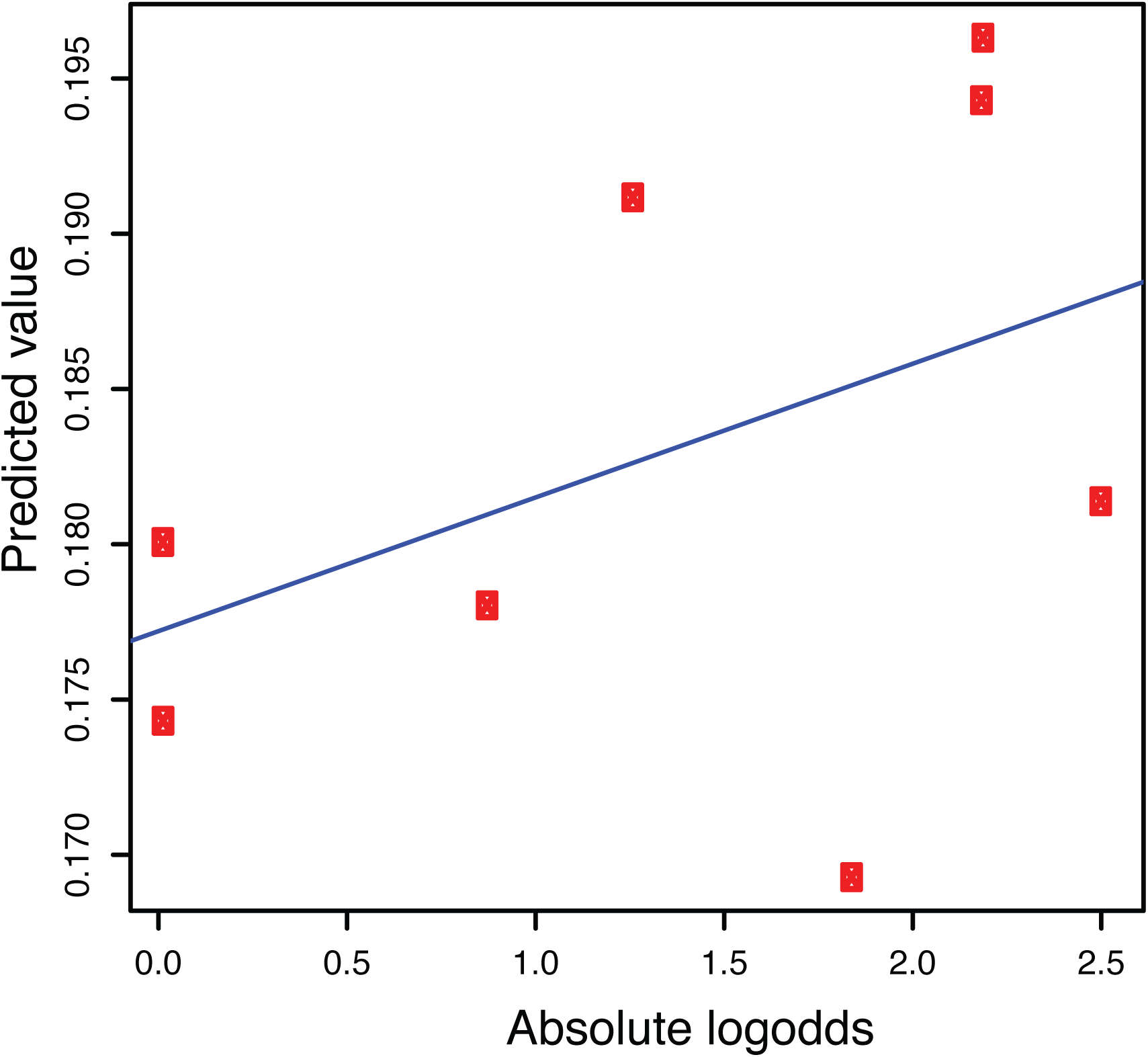
Predicted probability for emVar and non-emVAR versus absolute log2 odds from luciferase assay

