## Supplementary information for "GRAM: A generalized model to predict the molecular effect of a non-coding variant in a cell-type specific manner"

**S1** table Predictive performance of different feature sets, including cell-line specific ChIP-Seq TF binding scores and SELEX TF binding scores, using Lasso, SVM and Random Forest

| **MSE** | **Lasso 1se** | **SVM** | **Random Forest** |
| --- | --- | --- | --- |
| ChIPseq+SELEX | 0.106 | 0.105 | 0.102 |
| ChIPseq | 0.124 | 0.116 | 0.130 |
| SELEX | 0.111 | 0.108 | 0.107 |

S2 table Primers for 14 regions cloning in K562

| **Element Number** | **Forward Primer** | **Reverse Primer** |
| --- | --- | --- |
| 1 | GGGGACAACTTTGTACAAAAAAGTTGGCACCGGTACACGAAGGCTGGG | GGGGACAACTTTGTACAAGAAAGTTGGCAACTGCTGTAGCCCACCA |
| 2 | GGGGACAACTTTGTACAAAAAAGTTGGCACCGGCAGTTGGAAAGGGC | GGGGACAACTTTGTACAAGAAAGTTGGCAAGTCCTCAGGAACCGGC |
| 3 | GGGGACAACTTTGTACAAAAAAGTTGGCACCCGTGATCAACCAAAATCACCTCA | GGGGACAACTTTGTACAAGAAAGTTGGCAATCCTCTGCTCTACGGAATGGA |
| 4 | GGGGACAACTTTGTACAAAAAAGTTGGCACCTATCCGGAAGGGGTGGAACC | GGGGACAACTTTGTACAAGAAAGTTGGCAACCCCAGGTCACTCAGACCAC |
| 5 | GGGGACAACTTTGTACAAAAAAGTTGGCACCTAAGCCCAGTCCTCAGC | GGGGACAACTTTGTACAAGAAAGTTGGCAAAAGGAAGACTAATTGGGTCCC |
| 6 | GGGGACAACTTTGTACAAAAAAGTTGGCACCAGAACTAGCAGCGGAGAC | GGGGACAACTTTGTACAAGAAAGTTGGCAATCCTTCTTGCATGGTCGG |
| 7 | GGGGACAACTTTGTACAAAAAAGTTGGCACCTCCCCTGCTTTTCGATCTCTC | GGGGACAACTTTGTACAAGAAAGTTGGCAACTTTAGCCGCCCCTTTCCA |
| 8 | GGGGACAACTTTGTACAAAAAAGTTGGCACCCCTTTTCTAATCACCATTTCCGATA | GGGGACAACTTTGTACAAGAAAGTTGGCAATAACAGGGGCAATGCACTTT |
| 9 | GGGGACAACTTTGTACAAAAAAGTTGGCACCTGTTTTGCTTTTAACCTGATGAT | GGGGACAACTTTGTACAAGAAAGTTGGCAAAGGAGGTGGGGTGGCT |
| 10 | GGGGACAACTTTGTACAAAAAAGTTGGCACCCTTTCACGCAAAACCTGCTCA | GGGGACAACTTTGTACAAGAAAGTTGGCAAAACCTTTTTCCTTGAACCTTAGAC |
| 11 | GGGGACAACTTTGTACAAAAAAGTTGGCACCGGTCTTGACGCTGGCC | GGGGACAACTTTGTACAAGAAAGTTGGCAATGCTGGGAGAAACCGAT |
| 12 | GGGGACAACTTTGTACAAAAAAGTTGGCACCGGCTTCACTAACCTTAAATTCTAAA | GGGGACAACTTTGTACAAGAAAGTTGGCAATGCCCTTAAACAAGATGGC |
| 13 | GGGGACAACTTTGTACAAAAAAGTTGGCACCGTTCCCCTTCTGTCTCAGG | GGGGACAACTTTGTACAAGAAAGTTGGCAATTCATTCAGGGGCTCCC |
| 14 | GGGGACAACTTTGTACAAAAAAGTTGGCACCGACCCCACCCCTTCCC | GGGGACAACTTTGTACAAGAAAGTTGGCAATTTTGGCAGGAGCAGGA |

**Fig. S1** PRC curve for regulatory activity prediction

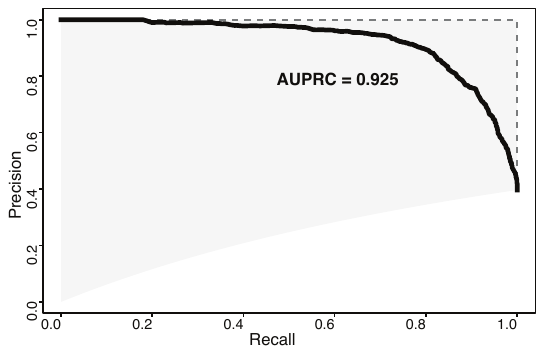

**Fig. S2** Principal component analysis using Vodds for three cell lines: GM12878, GM19239 and HepG2

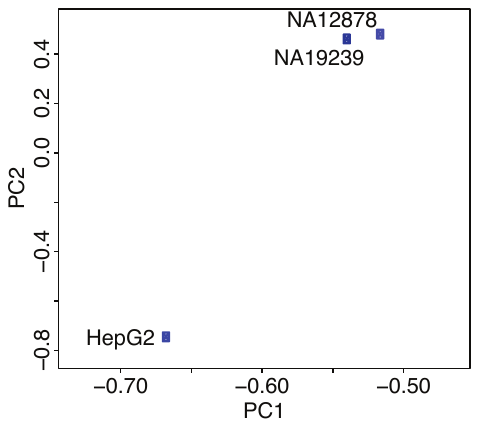

**Fig. S3** Distribution of Vodds score for GM12878. The high and low variable cell specificity class are defined by the top and bottom quantile.

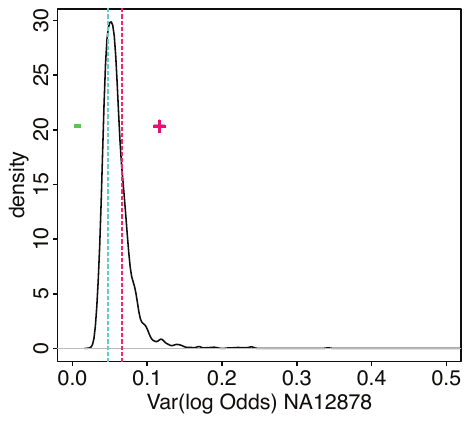

**Fig. S4** The prediction of cell type modifier score using TF binding profiles.

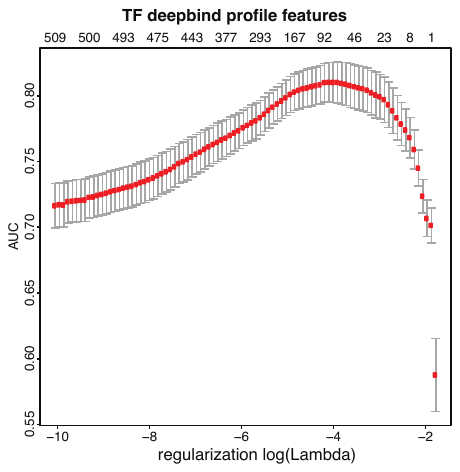

**Fig. S5** Predicted probability for emVar and non-emVAR versus absolute log2 odds from luciferase assay

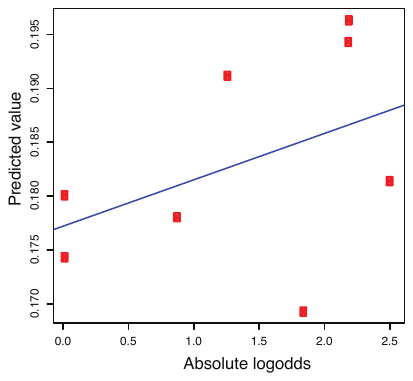
